# Structural basis for long-chain isoprenoids synthesis by *cis*-prenyltransferases

**DOI:** 10.1101/2021.10.21.465316

**Authors:** Moshe Giladi, Michal Lisnyansky Bar-El, Pavla Vaňková, Alisa Ferofontov, Emelia Melvin, Daniel Kavan, Boris Redko, Elvira Haimov, Reuven Wiener, Petr Man, Yoni Haitin

## Abstract

Isoprenoids are the largest group of natural products, found in all living organisms and play an essential role in numerous cellular processes. These compounds are synthesized by prenyltransferases, catalyzing the condensation reaction between an allylic diphosphate primer and a variable number of isopentenyl diphosphate (C_5_) units. This superfamily of enzymes can be subdivided into *trans*- or *cis*-prenyltransferases according to the stereoisomerism of the product. The *cis* branch can be further classified according to product length. While the active site volume was suggested to determine the final length in enzymes synthesizing short- and medium-chain products (up to C_60_), long-chain enzymes (up to C_120_) and rubber synthases (>C_10,000_) fail to conform to this paradigm. Here, to resolve the structural basis for long-chain isoprenoid synthesis, we focused on the human *cis*-prenyltransferase complex (h*cis*-PT). This enzyme, peripheral to the endoplasmic reticulum membrane, produces the precursor for dolichol phosphate, a membrane residing glycosyl carrier. In line with its crucial role in the cellular protein glycosylation machinery, disease-causing mutations in h*cis*-PT were shown to result in a wide spectrum of clinical phenotypes. The crystallographic structures of h*cis*-PT in four different substrate/product-bound conformations revealed an outlet enabling product elongation into the bulk solvent. Moreover, hydrogen-deuterium exchange mass spectrometry analysis in solution showed that the hydrophobic active site core is flanked by dynamic regions consistent with separate inlet and outlet orifices. Finally, using a fluorescent substrate analog and a fluorescently-labeled lipid nanodiscs, we show that product elongation and membrane association are closely correlated. Together, our results support directional product synthesis in long-chain enzymes and rubber synthases, with a distinct substrate inlet and product outlet, allowing direct membrane insertion of the elongating isoprenoid during catalysis. This mechanism uncouples active site volume from product length and circumvents the need to expulse hydrophobic product into a polar environment prior to membrane insertion.

## Introduction

Isoprenoids are a large group of natural products, abundant across all kingdoms of life.^1^ These compounds are crucial for numerous cellular processes, such as the biosynthesis of cholesterol, steroid hormones, visual pigments, and moieties for post-translational protein modifications.^2^ Isoprenoids are synthesized by prenyltransferases, a group of enzymes that catalyze the condensation reaction between an allylic diphosphate primer and an isopentenyl diphosphate (IPP, C_5_) building block.^3,4^ In accordance with the stereoisomerism of the double bond formed during the reaction, these enzymes are subdivided into *cis*- and *trans*-prenyltransferases. Adding further complexity to the system, *cis*-prenyltransferases are also classified into four major classes according to the length of the final product. These classes include short-chain (up to C_20_), medium-chain (up to C_60_), long-chain (up to C_120_) and rubber synthase (>C_10,000_).^5,6^ In addition to the difference in product chain length, *cis*-prenyltransferases differ in their subunit composition. Specifically, while short- and medium-chain producing enzymes are homodimeric, enzyme belonging to the long-chain and rubber synthase classes are heteromeric.^7^

Despite the huge variance in product chain length and subunit composition, all four *cis*-prenyltransferase classes share an overall conserved structure and catalytic mechanism.^5,6,8^ The active site is divided into two substrate bindings sites termed S_1_ and S_2_ (Figure 1A). The allylic diphosphate primer binds at S_1_, while IPP binds at S_2_. Once both sites are occupied by substrates, with a Mg^2+^ ion serving as an essential co-factor, the diphosphate group of the allylic primer is hydrolyzed, leaving behind a carbocation intermediate that undergoes condensation with IPP. Following the binding of a new IPP molecule at S_2_ the cycle can be repeated until a final chain length is reached and the product dissociates. Previously, structure-function studies of short- and medium-chain enzymes suggested that the final chain length is directly related to the volume of S_1_.^9–11^ However, the recently determined crystal structures of the long-chain human *cis*-prenyltransferase complex (h*cis*-PT) challenges this view (Figure. 1A).^12,13^ Specifically, the volume of S_1_ in the long-chain h*cis*-PT was surprisingly found to be smaller than that of medium-chain orthologs, making it clearly incompatible with long products. These observations accentuate our missing mechanistic understating of long-chain isoprenoids and rubber synthesis.

**Figure 1.**
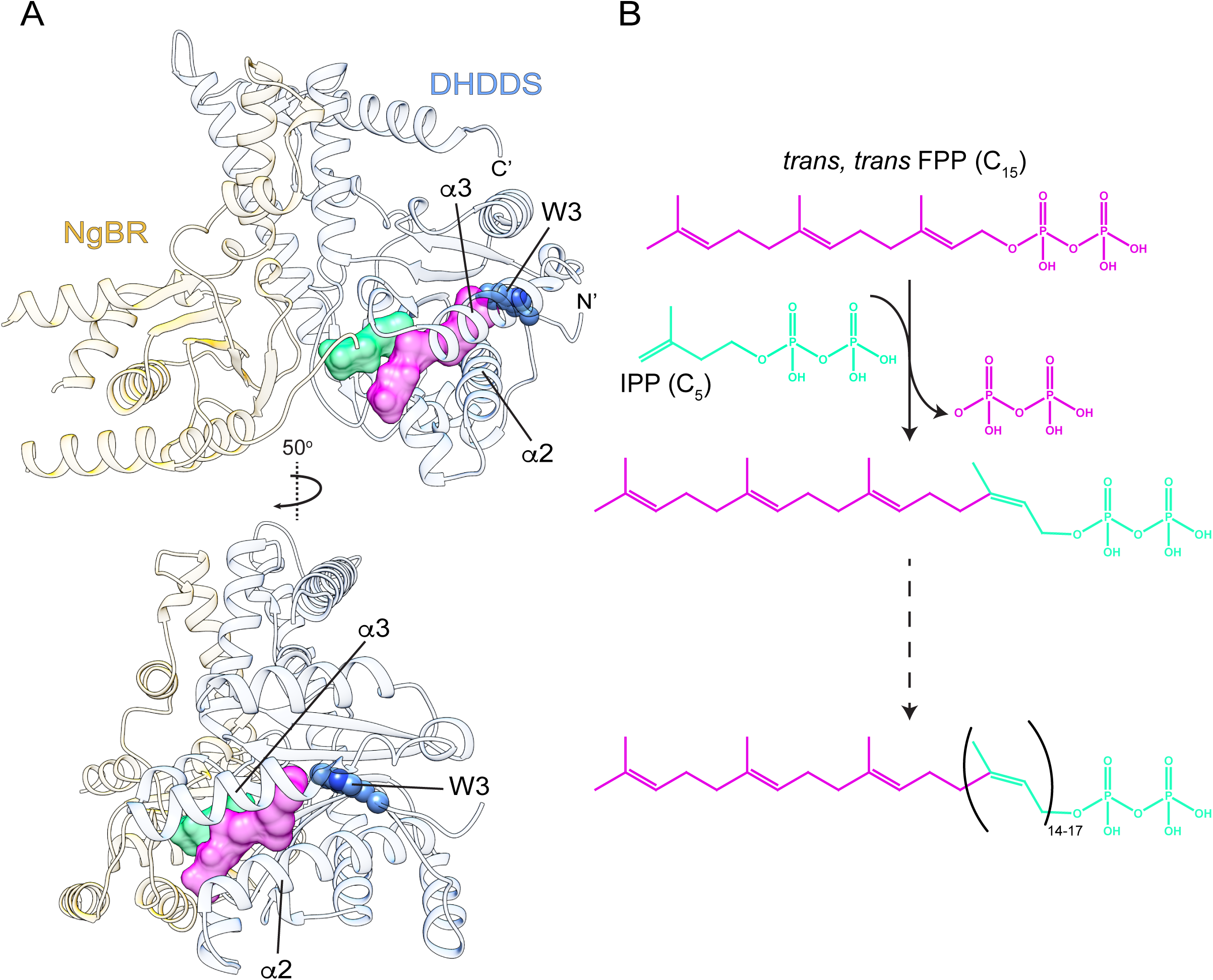
Overview of h*cis*-PT structure and reaction scheme. (A) Cartoon representation of a single DHDDS-NgBR heterodimer in complex with FPP (PDB 6Z1N). DHDDS and NgBR are colored blue and yellow, respectively. The pink and green surfaces represent the S_1_ and S_2_ sites, respectively. W3, at the DHDDS N-terminus, is shown as spheres. (B) The condensation reaction scheme. At the first cycle, the allylic diphosphate primer, FPP (C_15_, pink) undergoes a condensation with IPP (C_5_, green) to produce GGPP (C_20_). The cycle repeats with further condensations (14-17) of the allylic diphosphate at S_1_, ultimately leading to a final product length of C_85-100_.

h*cis*-PT plays an important role in protein N-glycosylation and thus, is ubiquitously expressed in every cell type.^14^ This heteromeric enzyme is composed of two types of subunits: a catalytic subunit termed dehydrodolichyl diphosphate synthase (DHDDS) and an auxiliary quiescent subunit termed Nogo B-receptor (NgBR) (Figure 1A).^15^ In line with its important biological roles, mutations in both subunits of h*cis*-PT were identified as causing several human diseases with ocular^16,17^ and neurological^18–21^ manifestations, among others.^22,23^ Physiologically, h*cis*-PT is localized to the endoplasmic reticulum (ER) membrane via an N-terminal transmembrane domain of NgBR, where it synthesizes the precursor for dolichol-phosphate (Dol-P) by consecutive condensations of IPP onto the allylic diphosphate primer farnesyl diphosphate (FPP, C_15_) (Figure 1B).^7,15^ Dol-P serves as a lipidic glycosyl carrier for N-glycosylation, residing in the ER membrane.^14,24^ The structures of the soluble regions of h*cis*-PT (sh*cis*-PT), devoid of the transmembrane region of NgBR,^12,13^ revealed that h*cis*-PT is a heterotetramer, formed by two DHDDS-NgBR heterodimers (Figure S1). DHDDS exhibits the common *cis*-prenyltransferase fold, composed of 7 α-helices (α1-7) and 6 β-strands (βA-F), flanked by an N-terminal helix (αN) and a unique C-terminal helix-turn-helix motif enabling heterotetramerization (Figure S1). Conversely, while NgBR exhibits significant sequence similarity with *cis*-prenyltransferases, its structure revealed a pseudo *cis*-prenyltransferase fold, composed of 6 α-helices (α1-6) and 5 β-strands (βA-E) (Figure S1).^25^ Although NgBR exhibits no endogenous *cis*-prenyltransferase activity, it accelerates that activity of DHDDS ~ 400-fold by direct interaction of the NgBR C-terminal tail with active site residues and substrates.^12,26^

In all *cis*-prenyltransferases, the hydrophobic pocket of S_1_ is surrounded by 2 helices (α2, α3) and 4 β-strands (βA, βB, βE, βF). However, the structural elements forming the active site “bottom”, juxtaposing the pyrophosphate binding region (Figure 1A), differ between the different classes. In short- and medium-chain enzymes, the bottom consists of rigidly and tightly packed bulky hydrophobic residues, leaving the active site inlet as the only orifice to the bulk solvent. Indeed, mutations of these bulky residues at the bottom of the active site change the available volume for product elongation, altering product length.^27^ In contrast, long-chain enzymes harbor a unique N-terminal tail forming the active site bottom. Specifically, an insertion in α3 leads to active site widening, allowing the N-terminal tail of DHDDS to snake into the active site, with the conserved W3 strategically situated as a stopcock preventing exposure of the hydrophobic pocket to the bulk solution (Figure 1A) by forming hydrophobic interactions with F55 (α2), F101 (α3), and V152 (βD).

Here, to elucidate the structural basis for long-chain isoprenoid synthesis by h*cis*-PT, we combined X-ray crystallography, hydrogen-deuterium exchange mass spectrometry (HDX-MS) and fluorescence spectroscopy. By determining multiple crystal structures of sh*cis*-PT in the presence of reactive and non-reactive substrate analogs, we halted catalysis and captured the enzyme in specific predefined states along its catalytic cycle. Strikingly, following a single condensation reaction, the N-terminus of DHDDS translocates, resulting in opening of a novel outlet through which the C_20_ product extends towards the bulk solvent. In line with the crystal structures, HDX-MS analysis of apo sh*cis*-PT reveals that the N-terminus of DHDDS is highly dynamic in solution, supporting the propensity of the novel outlet to form. Finally, fluorescence resonance energy transfer (FRET) measurements between a fluorescent FPP analog and fluorescently labeled lipid nanodiscs indicate direct product elongation into the lipid bilayer. Together, our results depict a product elongation mechanism involving its dynamic interplay with the N-terminal tail of DHDDS, where expulsion of W3 results in the formation of an outlet through which product immersion into the adjacent lipid bilayer can be achieved, circumventing the need for housing long-chain products. Moreover, this mechanism may also explain the activity enhancement of h*cis*-PT observed in the presence of detergents or lipids,^13,28^ as well as the molecular basis for natural rubber synthesis.^6^

## Results

### Crystallographic analysis of sh*cis*-PT along the first condensation reaction

Previously, the catalytic mechanism facilitating the condensation reaction by *cis*-prenyltransferases was extensively studied, with residues involved in the reaction mechanism shown to be highly conserved among the different classes.^5–8,10^ However, the molecular mechanism enabling the accommodation of long-chain products remains to be resolved. Hence, in order to dissect the conformational states along the condensation reaction, we analyzed the conformational transitions associated with the first catalytic cycle using reactive and non-reactive substrate analogs (Figure 2, Table 1). To this end, we purified sh*cis*-PT_cryst_, consisting of full-length DHDDS (residues 1-333) and the pseudo *cis*-prenyltransferase domain of NgBR (residues 73-293) lacking residues 167-175, followed by crystallization and structure determination in the presence of different substrates combinations. As sh*cis*-PT produces product population ranging in lengths (Figure 1B), this approach favors the formation of specific and highly homogeneous enzyme-substrate complexes, facilitating our structural investigations. In all four structures determined here, the asymmetric unit of the crystal consists of a single heterodimer. The heterotetramer is formed via a dimerization interface between adjacent asymmetric units, mainly consisting of the C-terminus of DHDDS (Figure S1).

**Figure 2.**
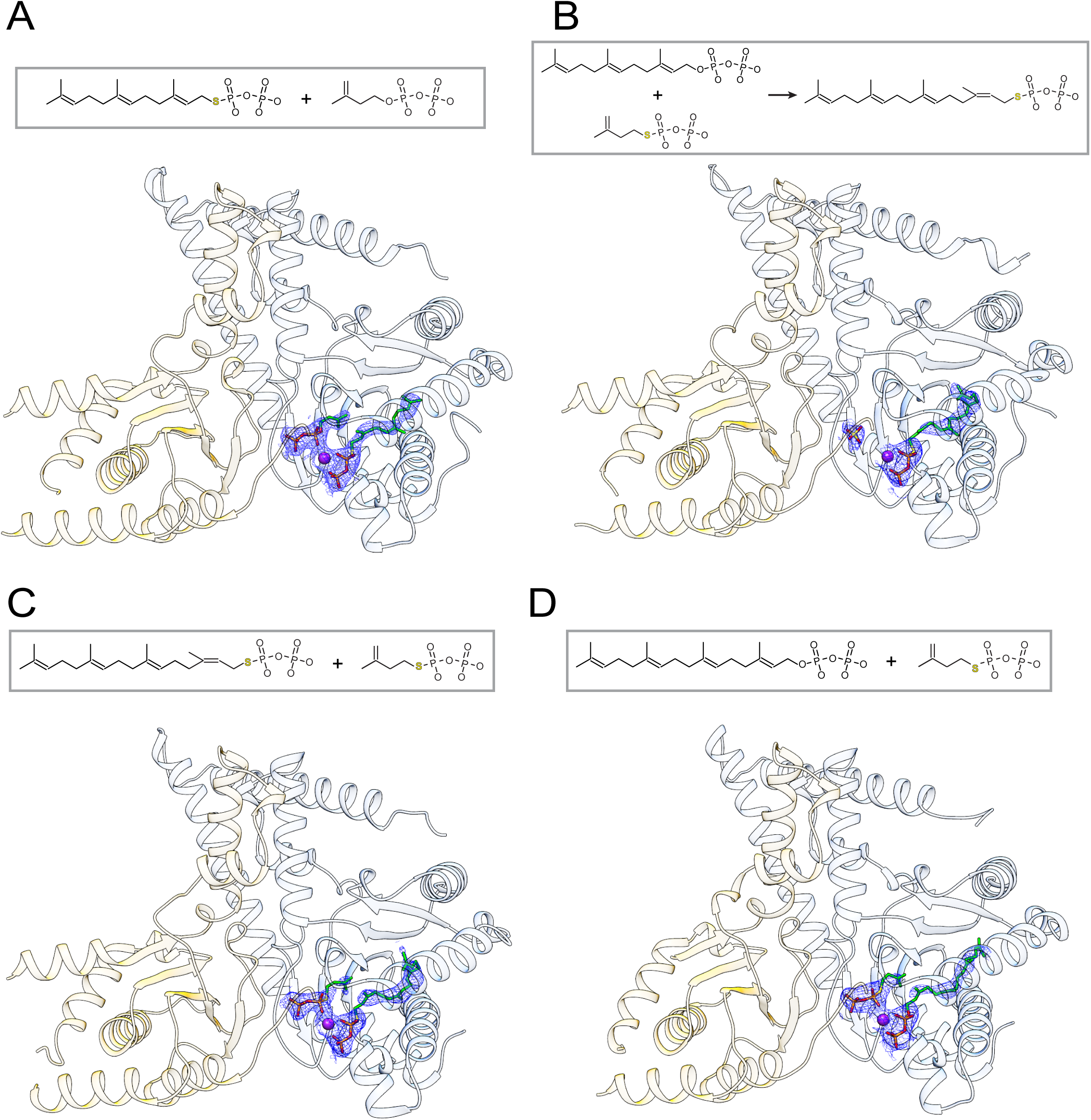
Structural analysis of sh*cis*-PT_crystal_ along the first condensation cycle. (A-D) Cartoon representation of a single DHDDS-NgBR heterodimer in complex with FsPP and IPP (A), GGsPP and sulphate (B), GGsPP and IsPP (C), or GGPP and IsPP (D). DHDDS and NgBR are colored blue and yellow, respectively. The bound substrates and/or products are shown as sticks, the Mg^2+^ ion is shown as purple sphere, and a composite omit map is shown as a blue mesh (contoured at σ = 1).

**Table 1.**
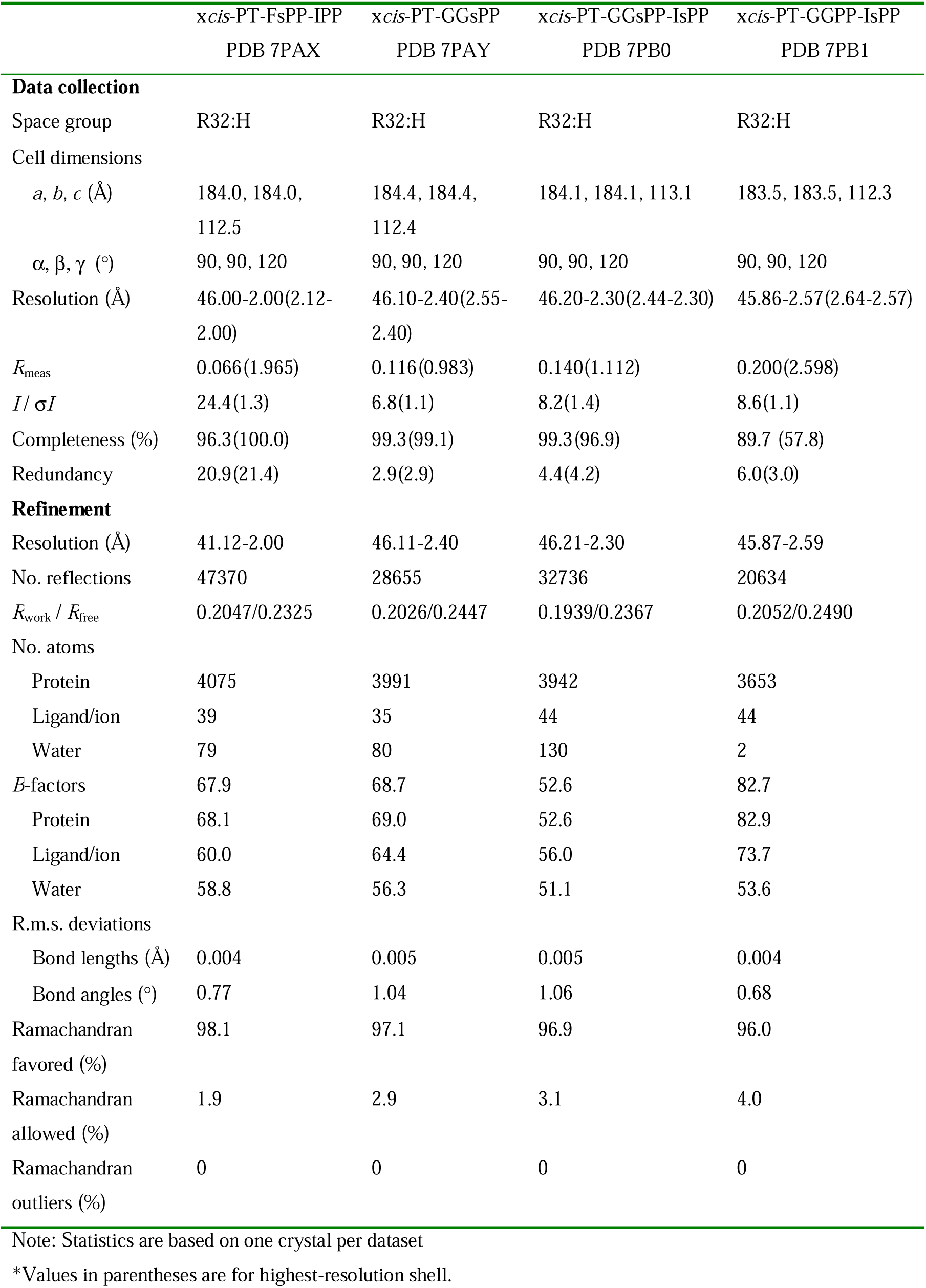
Data collection and refinement statistics.

To isolate the initial substrate-bound state, where FPP and IPP occupy S_1_ and S_2_, respectively, we crystallized sh*cis*-PT_cryst_ in the presence of Mg^2+^, farnesyl-thiodiphosphate (FsPP) and IPP.^29,30^ The replacement of the hydrolysable oxygen with a sulfur atom in FsPP results in a poorly reactive substrate compared with FPP (Figure 2A), allowing us to capture the pre-condensation state. Next, to resolve the enzyme harboring the product of the first condensation reaction, we crystallized the complex in the presence of Mg^2+^, FPP and isopentenyl-thiodiphosphate (IsPP) (Figure 3B).^30^ The condensation of FPP with IsPP results in the formation of geranylgeranyl-thiodiphosphate (GGsPP), which is recalcitrant for further condensations, due to the same mechanism as in the case of FsPP. Serendipitously, we were able to obtain two different forms of this GGsPP-bound enzyme, one containing a sulphate ion (Figure 3B) and the other IsPP (Figure 3C), bound at S_2_. Finally, we crystallized sh*cis*-PT_cryst_ in the presence of geranylgeranyl-diphosphate (GGPP) and IsPP as initial substrates (Figure 3D), with the goal of obtaining a structure corresponding to sh*cis*-PT_cryst_ in complex with the second condensation product farnesylgeranyl diphosphate (C_25_). However, the protein only crystallized in the pre-condensation state, with the substrates GGPP and IsPP bound at S_1_ and S_2_, respectively.

**Figure 3.**
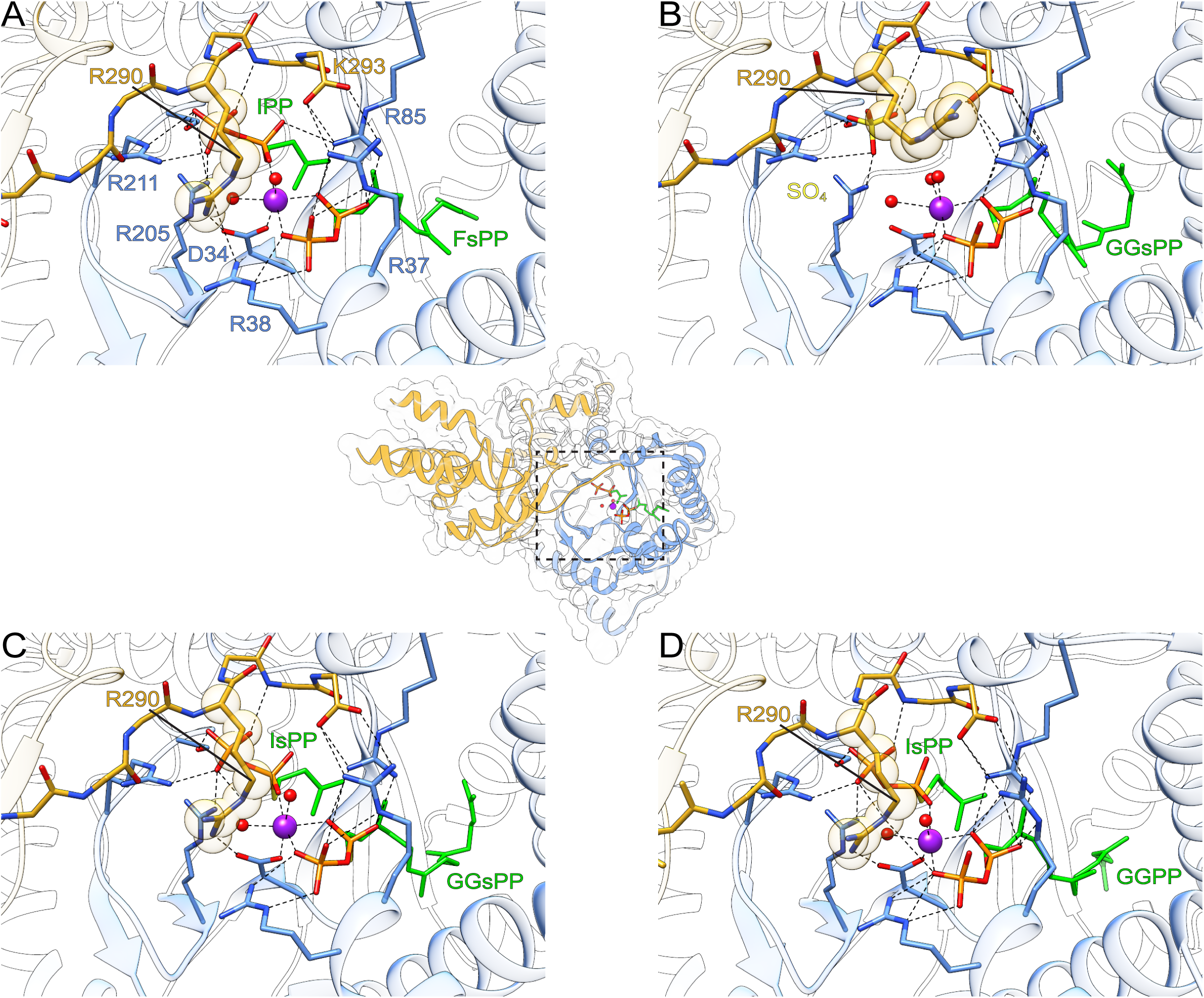
Structural organization of the active site inlet. (A-D) Zoom perspective of the S_1_ and S_2_ inlet, framed by a dashed rectangle on the DHDDS-NgBR heterodimer structure at the center. Mg^2+^ and water molecules are shown as purple and red spheres, respectively. Coordinating residues from DHDDS (blue), the C-terminus of NgBR (yellow) and substrates and/or products are shown as sticks. The structures in complex with FsPP and IPP (A), GGsPP and sulphate (B), GGsPP and IsPP (C) or GGPP and IsPP (D) are shown. Polar interactions are highlighted as dashed black lines, and R290 is highlighted by spheres.

### Substrate coordination at S_1_ and S_2_

Previously, the structure of sh*cis*-PT_cryst_ was determined in the presence of FPP at S_1_ and a phosphate group at S_2_^12^ or in a state where IPP occupies both sites.^13^ In contrast, the structures reported here represent native states, in which both sites are occupied by their respective substrates. Thus, we first focused on the organization of the superficial polar region, involved in pyrophosphate binding, along the reaction cycle (Figure 3). At S_1_, the pyrophosphate group of FsPP (Figure 3A), GGsPP (Figure 3B, C) or GGPP (Figure 3D) interacts with a Mg^2+^ ion and is also stabilized by interactions with R37, R38 and R85 from DHDDS. At S_2_, the pyrophosphate group of IPP (Figure 3A), the sulphate ion (Figure 3B) or the pyrophosphate of IsPP (Figure 3C,D) are stabilized by R205, R211, S213 from DHDDS and the backbone nitrogen of G292 from NgBR. In all the structures, the NgBR C-terminal carboxylate (K293) interacts with R37 and R85 of DHDDS, involved in substrate coordination at S_1_ (Figure 3).

The structure with bound FsPP and IPP (Figure 2A, 3A) reveals that the simultaneous occupancy of S_1_ and S_2_ does not alter the conformation of the substrates, compared with structures obtained in complex with either substrate alone (Figure S2). However, the presence of IPP (Figure 3A) or IsPP (Figure 3C,D), rather than sulphate (Figure 3B) or phosphate (Figure 1A), at S_2_ results in two main changes. First, the orientation of R290 from NgBR is altered (Figure 3A,C,D *vs*. Figure 3B). Specifically, while R290 does not interact with the sulphate moiety, it directly interacts with D34 and the substrates in the IPP or IsPP bound states. Intriguingly, underscoring the potential importance of these interactions for catalysis, the R290H mutation reduces catalytic activity *in vitro* and results in a congenital glycosylation disorder.^12,22,28^ Second, while Mg^2+^ is octahedrally coordinated via six oxygen atoms, contributed by the conserved D34, the pyrophosphate of FPP and water molecules in all the structures, one water molecule is replaced by an oxygen from the pyrophosphate group of IPP or IsPP (Figure 3A,C,D). Previously, it was proposed that after FPP binding to S_1_, a preformed Mg^2+^-IPP complex binds to S_2_, followed by translocation of the Mg^2+^ ion closer to S_1_, enabling the facilitation of pyrophosphate hydrolysis.^5^ Thus, the structures containing an allylic diphosphate at S_1_ and IPP or IsPP at S_2_, in which the Mg^2+^ ion is coordinated by the pyrophosphate moieties from both sites, likely represent an intermediate in Mg^2+^ translocation.

### An interplay between chain elongation and DHDDS N-terminus conformation

Previous crystallographic investigations of sh*cis*-PT_cryst_ revealed that the N-terminus of DHDDS snakes into the active site, blocking a potential outlet for the elongating product (Figure 1A).^12,13^ In the presence of FsPP and IPP (Figure 4A), the conformations of the farnesyl (C_15_) moiety and the N-terminus of DHDDS are similar to those previously observed in the FPP-bound state (Figure 1A, S2). Surprisingly, the structures harboring GGsPP or GGPP, following the first condensation reaction (Figure 2B,C,D), demonstrate distinct conformations of the geranylgeranyl (C_20_) moiety and the N-terminus of DHDDS (Figure 4B,C,D). Specifically, unlike the structure with FPP, both GGsPP-bound structures show bending of the geranylgeranyl moiety at C_11_, resulting from interaction of the terminal carbon (C_20_) with F154 (βD) within the hydrophobic pocket (Figure 4B,C). Interestingly, comparison of the two GGsPP-bound structures reveals that the N-terminus of DHDDS (up to G7) is only resolved in the absence of IsPP (Figure 4E,F), pointing towards the dynamic nature and consistent with the high B-factors of this region. Thus, the GGsPP-IsPP bound form reveals a conformation in which the hydrophobic pocket is vacant of W3 and exposed to the bulk solvent. This conformation represents a state in which the enzyme is predisposed for product expulsion and subsequent elongation. In accordance, the GGPP-bound structure strikingly revealed an elongated conformation, where the terminal carbons are expulsed from the hydrophobic pocket and are exposed to the bulk solvent (Figure 4D,E,F). Collectively, the structures of sh*cis*-PT_cryst_ along the reaction cycle reveal that both the product and the DHDDS N-terminus are inherently flexible, allowing the formation of a novel outlet for streamlined synthesis of long-chain products.

**Figure 4.**
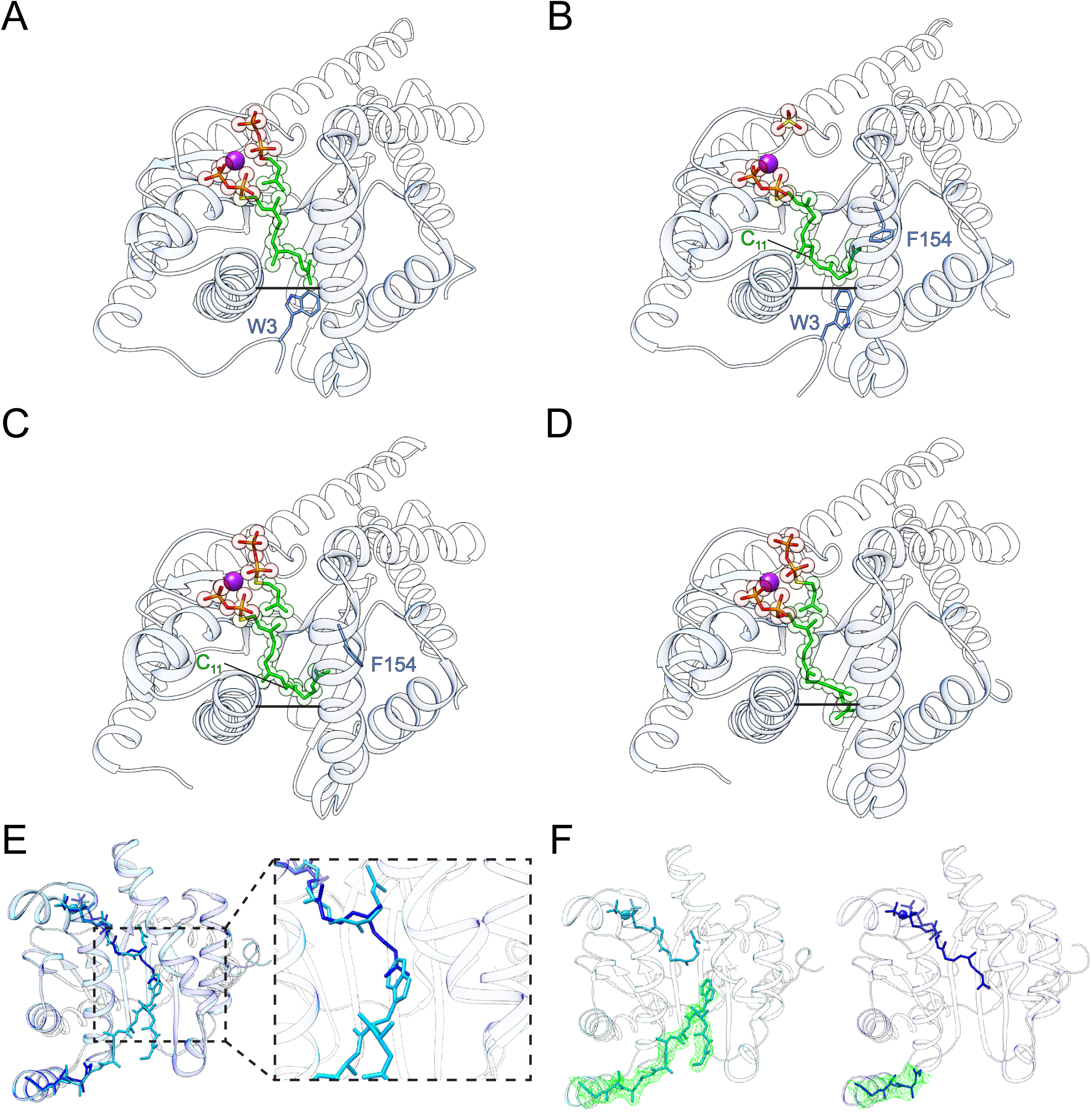
The interplay between the allylic diphosphate at S_1_ and DHDDS N-terminus conformations. (A-D) Cartoon representation of DHDDS in complex with FsPP and IPP (A), GGsPP and sulphate (B), GGsPP and IsPP (C), GGPP and IsPP (D) shown as sticks. W3, serving as a stopcock for the active site outlet, and F154, stabilizing the bent conformation of the C_20_ product within the active site, are shown as sticks. The level of the outlet is indicated as a solid black line between α2 and α3. (E) Superposition of sh*cis*-PT_cryst_ in complex with GGsPP (cyan) and GGPP-IsPP (blue). The N-terminus of DHDDS, GGsPP and GGPP are shown as sticks. (F) 2*F*_o_-*F*_c_ electron density maps of the N-terminus of DHDDS, contoured at σ = 1, are shown as green mesh for sh*cis*-PT_cryst_ in complex with GGsPP (left) and GGPP-IsPP (right). No electron density could be identified upstream of residue G7.

### HDX-MS analysis reveals that the N-terminus of DHDDS is highly dynamic in solution

To determine the conformational dynamics of the N-terminus, governing the formation propensity of the novel hydrophobic pocket outlet, we performed HDX-MS analysis of apo sh*cis*-PT. In this method, the protein sample is diluted in deuterated buffer solution, resulting in the exchange of backbone amide hydrogens with deuterium.^31,32^ The reaction is quenched at different time points, followed by proteolytic digestion and quantitation of deuterium incorporation within each peptide.^32^ Importantly, the exchange rate reflects the local fold and solvent accessibility of each protein region. Following proteolytic digestion, we were able to obtain a 100% and 98.7% coverage for DHDDS and NgBR, respectively (Figure S3).

Overall, sh*cis*-PT exhibits a characteristic HDX pattern for a soluble protein, where the core, secondary structure elements and buried interfacial regions exhibit low HDX (Figure 5A). Focusing on the active site, the HDX profile reveals three distinct regions displaying alternating HDX levels (Figure 5). First, the inlet, composed of the polar pyrophosphate binding regions of S_1_ and S_2_. The inlet (Figure 5A) includes α1, the N-terminal portions of α2 and α3 and the C-terminal tail of NgBR (Figure 5B), and encompasses several key positively charged residues (R37, R38, R85, R205, R211) involved in coordination of the pyrophosphate moieties (Figure 3). This region exhibits relatively high HDX, consistent with previous studies of medium-chain orthologs, suggesting local unfolding in the absence of substrates. Second, the hydrophobic pocket (Figure 5A), which includes the C-terminal portion of α2, the middle of α3, βA, βB, the C-terminal portion of βC, and βD (Figure 5B). This region exhibits the lowest HDX as expected from a rigid, well-structured and solvent-inaccessible segment. Finally, the outlet (Figure 5A), composed of the C-terminal portion of α3, the N-terminal portion of βC and the N-terminus of DHDDS (residues 2-11) (Figure 5B). This region exhibits higher HDX compared to the hydrophobic core. Importantly, the dynamics of the C-terminal portion of α3 are in agreement with its previously suggested role in active site widening and accommodation of long-chain products during catalysis. Moreover, the N-terminus of DHDDS exhibits notably high HDX throughout the experiment (Figure 5B), indicating that it is highly labile in solution. Together, the alternating pattern of solvent exposure within the active site supports directional product processing, with an inlet at the pyrophosphate binding region and an outlet at the distal hydrophobic pocket.

**Figure 5.**
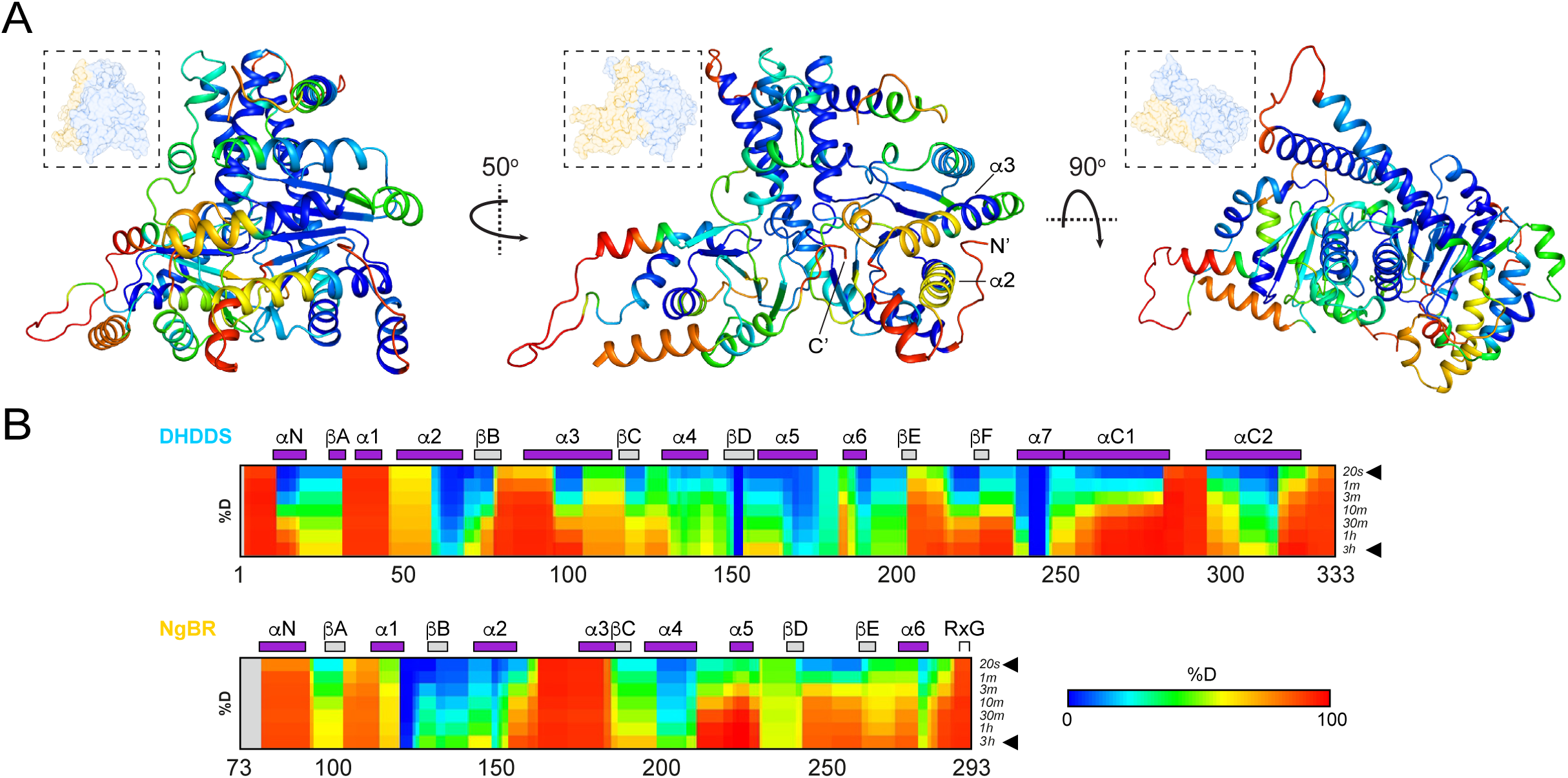
HDX-MS analysis of sh*cis*-PT. (A) Cartoon representations of a single DHDDS-NgBR heterodimer. Heat map coloring represents the HDX level following 20 sec incubation in D_2_O, with highest and lowest exchange levels indicated in red and blue, respectively. In each orientation, an inset provides a DHDDS (blue) and NgBR (yellow) heterodimer surface representation, for orientation purpose. The N-terminus, α2 and α3 of DHDDS, and the C-terminus of NgBR are indicated. (B) Deuteration levels at the indicated time points for DHDDS (upper panel) and NgBR (lower panel). The secondary structures elements are indicated above the heat maps for each protein.

### Use of a fluorescent FPP analog for structure-function investigations of sh*cis*-PT

Long-chain *cis*-prenyltransferases and rubber synthases are membrane delimited, and the presence of detergents or phospholipids was shown to dramatically enhance their catalytic activity.^6,13,28^ As our structural studies suggest product elongation through a novel outlet, possibly into the adjacent membrane, we sought to monitor the chemical environment and position of the elongating isoprene moiety. To this end, we utilized the fluorescent FPP analog (2E,6E)-8-O-(N-methyl-2-aminobenzoyl)-3,7-dimethyl-2,6-octandien-1-pyrophosphate (MANT-O-GPP),^33^ which was previously used to measure the activity of medium-chain enzymes.

To validate that MANT-O-GPP is a *bona fide* substrate for sh*cis*-PT, we used a radioactivity-based enzyme kinetics assay (Figure 6A), revealing that MANT-O-GPP exhibits K_m_ = 0.14 ± 0.01 μM (n = 3), similar to that previously reported for FPP.^12^ Next, as the excitation profile of MANT-O-GPP closely resembles the emission profile of tryptophan (Figure 6B), we reasoned that FRET can be used to assess the binding and position of MANT-O-GPP at S_1_. Specifically, the fluorescent MANT moiety is expected to reside in sub-nanometer proximity with W3 (Figure 6C), within FRET distance.^34^ Indeed, by using tryptophan excitation (*F*_280_) in the presence of increasing [MANT-O-GPP], we observed an increase in FRET. This is reflected by the concomitant decrease in tryptophan (*F*_340_) and increase in MANT-O-GPP (*F*_420_) emissions (Figure 6D). Notably, no direct MANT-O-GPP fluorescence is observed by excitation with *F*_280_ in the absence of sh*cis*-PT (Figure 6D). Moreover, following incubation with constant [MANT-O-GPP] (10 μM) and increasing [FPP], we observed an FPP-dependent FRET reduction (IC_50_ = 10.5 ± 3.0 μM, n = 3). This is reflected by an increase in *F*_340_ and a reciprocal decrease in *F*_420_, indicating that MANT-O-GPP and FPP share a common site. Finally, as DHDDS and NgBR contain eight and three tryptophan residues, respectively, each possibly contributing to the FRET signal, we sought to dissect the relative contribution of W3. Therefore, we generated the W3L mutant, maintaining the hydrophobicity of this position while abolishing its fluorescence (Figure 6F). While the WT and W3L mutant exhibit MANT-O-GPP-dependent FRET (Figure 6F), with a nearly identical K_d_ for MANT-O-GPP (1.06 ± 0.04 μM *vs*. 0.94 ± 0.09 μM, P = 0.33, n = 3), the lack of tryptophan at position 3 results in a significant reduction in the *F*_420_/*F*_340_ ratio (2.99 ± 0.04 *vs*. 1.77 ± 0.05, P < 0.0001, n = 3) (Figure 6G). Thus, MANT-O-GPP resides in close proximity to W3, as FPP, at the outlet of S_1_.

**Figure 6.**
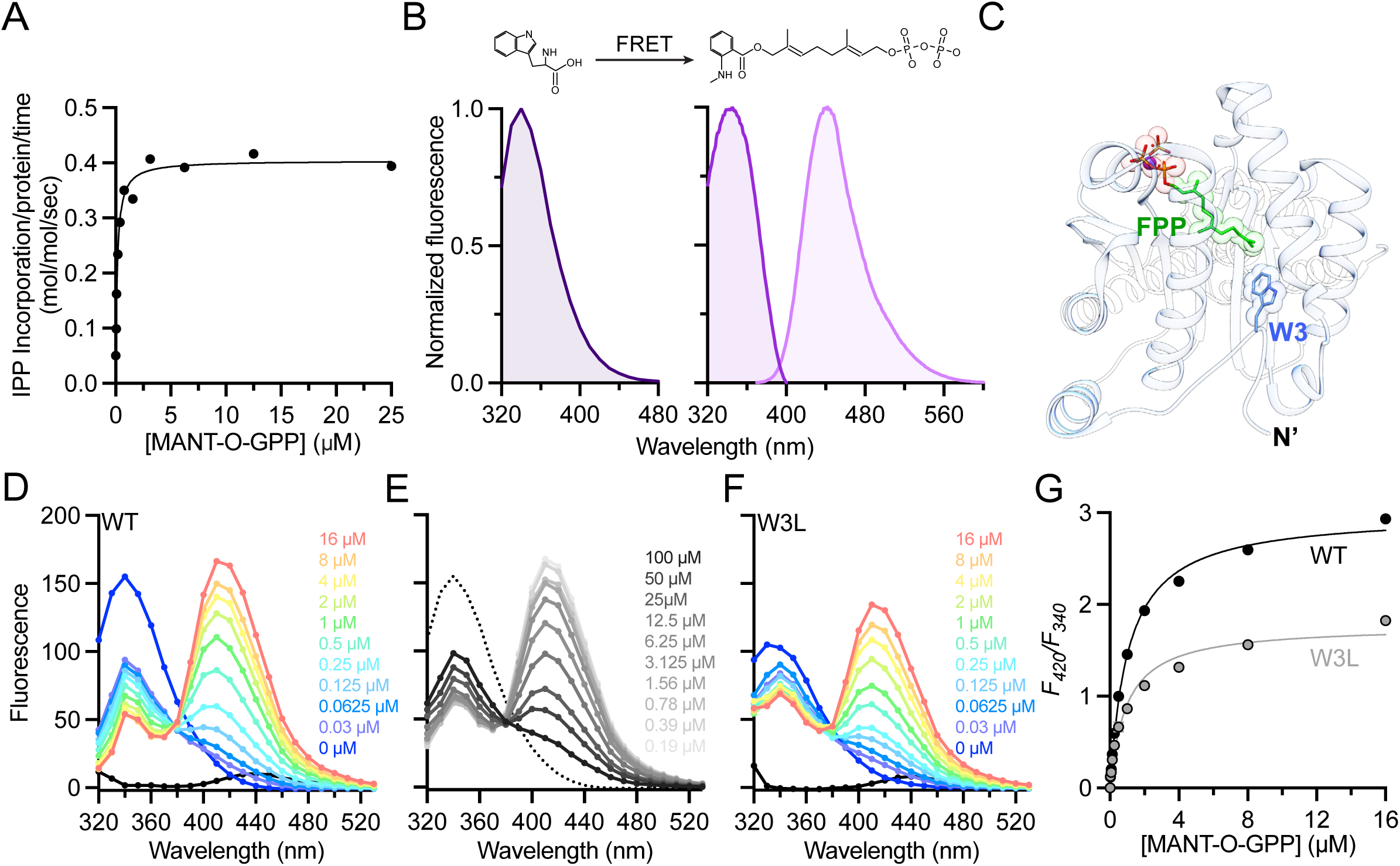
MANT-O-GPP binds at S_1_ and is a viable substrate of sh*cis*-PT. (A) Cartoon representation of sh*cis*-PT_cryst_ in complex with FPP (PDB 6Z1N), highlighting the close proximity between FPP (green sticks) and W3 (blue sticks). (B) Tryptophan emission spectrum (left panel) and MANT-O-GPP excitation (purple) and emission (pink) spectra (right panel). The chemical schemes of tryptophan and MANT-O-GPP are depicted. (C) MANT-O-GPP-dependent sh*cis*-PT activity in the presence of 100 μM IPP, measured as ^14^C-IPP incorporation. (D-F) Fluorescence emission spectra following excitation at *F*_280_ nm. Increasing concentrations of MANT-O-GPP resulted in concomitant decrease of *F*_340_ and increase of *F*_420_ in sh*cis*-PT-WT (D) or W3L mutant (F). Increasing FPP concentrations, in the presence of constant MANT-O-GPP (10 μM) resulted in concomitant increase of *F*_340_ and decrease of *F*_420_, indicating MANT-O-GPP displacement. (G) *F*_340_*/F*_420_ as a function of MANT-O-GPP concentration, measured using sh*cis*-PT-WT or W3L.

### sh*cis*-PT catalytic activity correlates with product-membrane interaction

To determine if the membrane bilayer can interact with, and possibly house, the elongating product, we first assessed the effect of azolectin nanodiscs on the catalytic activity of sh*cis*-PT. These measurements rely on the increased inherent MANT-O-GPP emission at *F*_420_ upon chain elongation.^33^ Consistent with the ability of phospholipids to enhance sh*cis*-PT activity, inclusion of nanodiscs at increasing nanodisc/enzyme ratios resulted in increased fluorescence amplitude and slope over the reaction time course (Figure 7A). Next, we measured FRET between MANT-O-GPP, serving as a donor, and azolectin nanodiscs containing 1% mol/mol 1,2-dioleoyl-sn-glycero-3-phosphoethanolamine-N-(7-nitro-2-1,3-benzoxadiazol-4-yl) (PE-NBD), a fluorescently labeled phospholipid, serving as an acceptor (Figure 7B). Following incubation of sh*cis*-PT with MANT-O-GPP, IPP and the fluorescent nanodiscs, we measured the fluorescence spectrum resulting from excitation of MANT-O-GPP (*F*_352_) (Figure 7B). In the absence of active catalysis (e.g. in the presence of EDTA or the absence of sh*cis*-PT or IPP), no fluorescence peak corresponding to PE-NBD emission is observed. However, inclusion of all components required for catalysis resulted in FRET between MANT-O-GPP and PE-NBD, as reflected by the emergence of a fluorescence peak at *F*_532_. These results indicate that chain elongation is a prerequisite for the FRET between the MANT and NBD moieties. Importantly, time-dependent measurements of the inherent MANT-O-GPP fluorescence and the MANT-NBD FRET were nearly identical (Figure 7D). To determine the minimal length required for product-membrane interaction, we limited the reaction to a single condensation, and monitored the FRET signal following incubation of sh*cis*-PT with MANT-O-GPP, fluorescent nanodiscs and IsPP in lieu of IPP (Figure 7E). This measurement revealed that even a single condensation is sufficient to give rise to FRET between the MANT and NBD moieties, although diminished. Together, these results support the notion that during long-chain isoprenoids synthesis by h*cis*-PT product elongation and its membrane interaction are interlaced.

**Figure 7.**
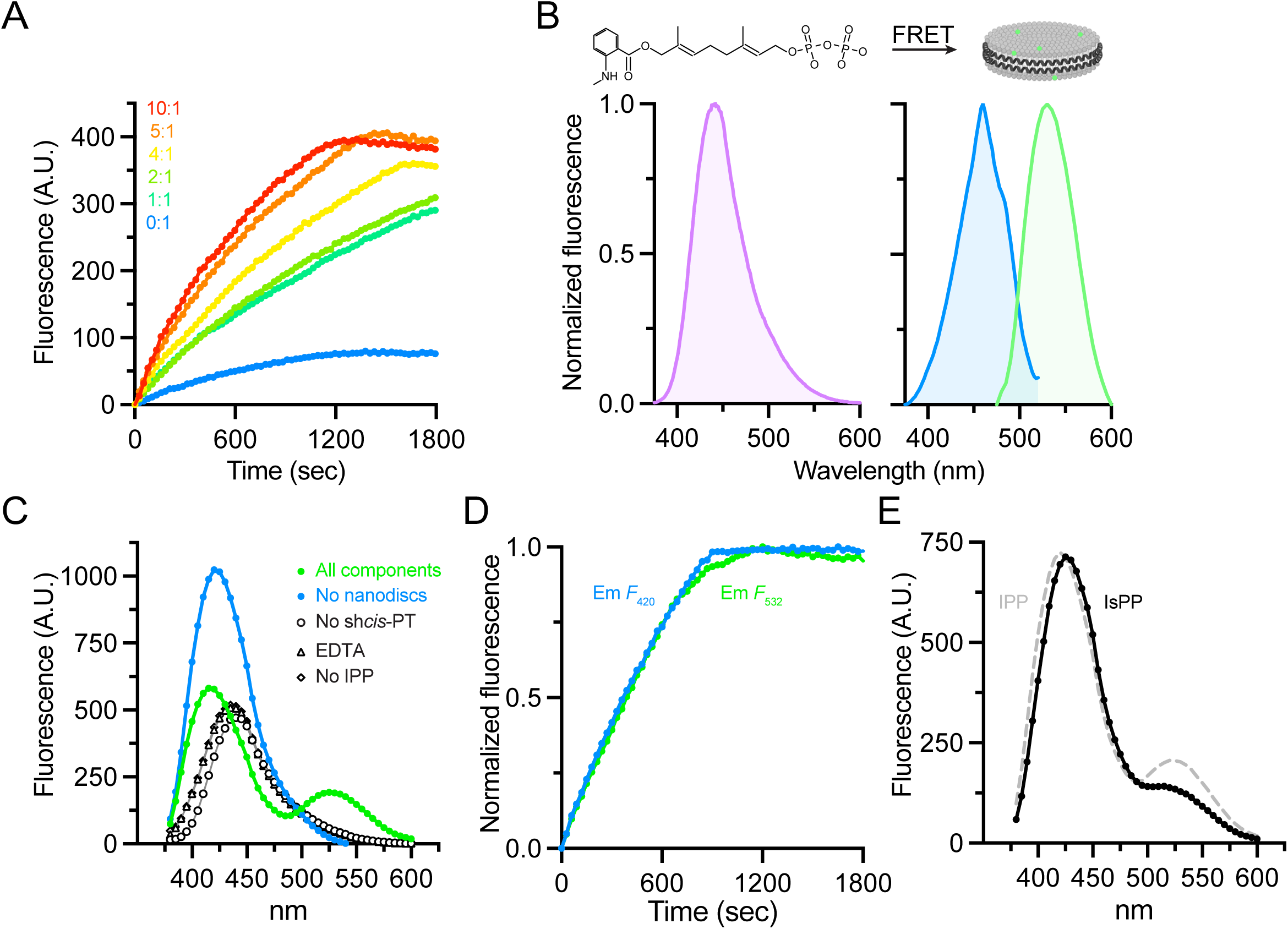
Monitoring sh*cis*-PT product-lipid interaction using FRET. (A) Time-dependent increase in MANT-O-GPP fluorescence (*F*_420_) in the presence of increasing MSP1D1E3 nanodiscs molar ratios. (B) MANT-O-GPP emission spectrum (left panel) and PE-NBD excitation (blue) and emission (green) spectra (right panel). The experimental scheme is depicted above spectra. (C) Fluorescence emission spectra following excitation at *F*_352_ nm. Incubation of sh*cis*-PT in the presence of MANT-O-GPP, IPP-Mg^2+^, and PE-NBD labeled nanodiscs (“all components”) resulted in a fluorescence peak corresponding to PE-NBD emission (*F*_532_) due to FRET (green). No FRET was observed upon omitting any of the reaction constituents. (D) MANT-O-GPP fluorescence (*F*_420_) and PE-NBD FRET (*F*_532_) exhibit nearly identical time-dependence. (E) Comparison of the fluorescence emission spectra following excitation at *F*_352_ nm and incubation of sh*cis*-PT in the presence of “all components” with IsPP instead of IPP (black). The emission spectra in the presence of IPP is shown as dashed grey curve.

## Discussion

Members of the *cis*-prenyltransferase family show high level of sequence conservation, translating into highly similar three-dimensional structures.^5,7,10^ Although sharing an identical catalytic mechanism, the final product length drastically varies between different *cis*-prenyltransferase sub-classes.^5,6^ While in short- and medium-chain enzymes the product length correlates with active site volume,^9–11^ recent structural analyses of the long-chain h*cis*-PT revealed that its active site is seemingly incompatible with such product lengths (Figure 1).^12,13^ The mismatch between the active site volume and product size is expected to aggravate in rubber synthases, producing extremely long (>C_10,000_) isoprenoids.^6^ Thus, products generated by the long-chain and rubber synthase sub-classes were hypothesized to elongate directly into neighboring cellular membranes, obviating the need for a large active site and implying a directional product expulsion via a previously unknown outlet during catalysis.^6,12^ Here, we used a hybrid approach to resolve the structural basis and determine the mechanism for long-chain isoprenoids synthesis.

Previously determined structures of sh*cis*-PT in complex with either FPP or IPP revealed that the hydrophobic pocket of DHDDS is occluded from the hydrophilic environment by the conserved W3 situated at the N-terminus of DHDDS (Figure 1A).^12,13^ The close proximity between FPP and W3 suggested that chain elongation should lead to translocation of the DHDDS N-terminus, thereby opening an alternative outlet through which the product can directly and directionally be expulsed. Thus, we crystallized sh*cis*-PT_cryst_ in the presence of different reactive and non-reactive substrates with a goal to capture the enzyme at different points along its catalytic cycle (Figure 2). The structures revealed that the active site experiences conformational plasticity along the catalytic cycle to enable long-chain isoprenoids synthesis.

Analysis of the different states captured (Figure 2) revealed that overall, the organization of the active site inlet, in which the condensation reaction takes place, is maintained (Figure 3). Nevertheless, we noticed that R290 of NgBR, a position previously identified as bearing a mutation (R290H) leading to a congenital glycosylation disorder,^22^ switches between two conformations in a substrate-dependent manner. Specifically, R290 only interacts with the substrates when both S_1_ and S_2_ are occupied (Figure 3). This conformational switch may support the previously suggested consecutive substrate binding model for *cis*-prenyltransferases, in which FPP binding to S_1_ precedes the binding of Mg^2+^-IPP to S_2_.^5^ In order to interact with the pyrophosphate at S_1_ and facilitate its hydrolysis, the Mg^2+^ ion must transverse between the sites. In the crystal structures resolved here, where both sites are occupied prior to the initiation of pyrophosphate hydrolysis, Mg^2+^ and R290 assume an intermediate position between the sites and interact with the pyrophosphates of both substrates. Thus, by providing charge compensation, it is possible that R290 relieves the Mg^2+^ from S_2_, allowing its translocation to S_1_.

Our structural analyses further revealed several distinct conformations of the enzyme-product complex (Figure 4). First, following the condensation and prior to the binding of an additional IPP molecule (Figure 4B), the C_20_ product is captured in a bent conformation (at C_11_), stabilized by hydrophobic interactions with F154 within the hydrophobic pocket. Here, the stopcock N-terminus of DHDDS maintains a closed conformation, as observed prior to the condensation reaction (Figure 1A, 4A). Next, when an additional substrate molecule occupies S_2_, the stopcock switches to an open state, as reflected by the lack of an identifiable electron density for the N-terminus of DHDDS (Figure 4C), forming a new outlet for further product elongation. Curiously, the bent conformation of the product molecule is maintained. This structural plasticity also occurs in solution, as supported by the high deuterium uptake of the DHDDS N-terminus throughout the HDX-MS experiment (Figure 5). Finally, the product assumes an elongated conformation, protruding through the new outlet (Figure 4D,E,F), keeping the stopcock at an open state. This structural organization is in sharp contrast to that previously observed in short- and medium-chain *cis*-prenyltransferases, where the hydrophobic pocket is tightly sealed by a rigid layer of bulky and hydrophobic residues.^35^ In addition, in these enzymes α3 is shorter and assumes a straight rather than a bent conformation, resulting in a rigid and narrow hydrophobic pocket.^5,6,9^ These structural differences preclude the opening of an outlet corresponding to that observed in sh*cis*-PT, limiting the product length to the active site dimensions.

In cells, the h*cis*-PT complex is an ER-resident enzyme, anchored to the membrane via the N-terminus of NgBR.^15^ In addition, the presence of hydrophobic environment was shown to dramatically stimulate enzymatic activity. However, this stimulation is abolished by mutations in hydrophobic residues located at the amphipathic αN of DHDDS, suggesting that interaction with the membrane is crucial for long-chain isoprenoids synthesis.^13^ Combined, these properties align with the proposal that the N-terminus of DHDDS can change its conformation to open the active site outlet, allowing direct product elongation into the adjacent membrane bilayer. Indeed, by monitoring the FRET signal between fluorescent substrate (Figure 6) and nanodiscs (Figure 7), we observed a close proximity between the enzymatic product and the membrane bilayer (Figure 7C). Moreover, the time course of chain elongation and emergence of the FRET signal were closely correlated (Figure 7D). The direct extension of the elongating product, resembling the passing of a thread through the eye of a needle, into the adjacent bilayer, solves two problems. First, as described above, it dismantles the relationship between product length and active site volume, keeping a constant portion of the product within the active site throughout the sequential condensations. Second, as product elongation results in its increased hydrophobicity, concomitant elongation and membrane insertion also serves to avoid the energetic barrier associated with hydrophobic product solvation.

Together, our results support a mechanism in which prior to the initial condensation reaction, the stopcock formed by the N-terminus of DHDDS occludes the active site outlet, shielding its hydrophobic interior from the bulk solvent (Figure 8A). Following the initial condensation, the stopcock opens and the product extends via the newly formed outlet, enabling the product to come in close proximity to the membrane and eventually to insert into it (Figure 8B). In principle, this mechanism alleviates any limitation on the final product length and can thus support natural rubber synthesis as well. Notably, when expressed in yeast cells, subunits of rubber synthase produce C_80-110_ products rather than the >C_10,000_ natural rubber produced when coupled with rubber particles.^36^ This suggests that long-chain enzymes and rubber synthases share a basic common synthetic mechanism, further modulated and fine-tuned by additional constituents.

**Figure 8.**
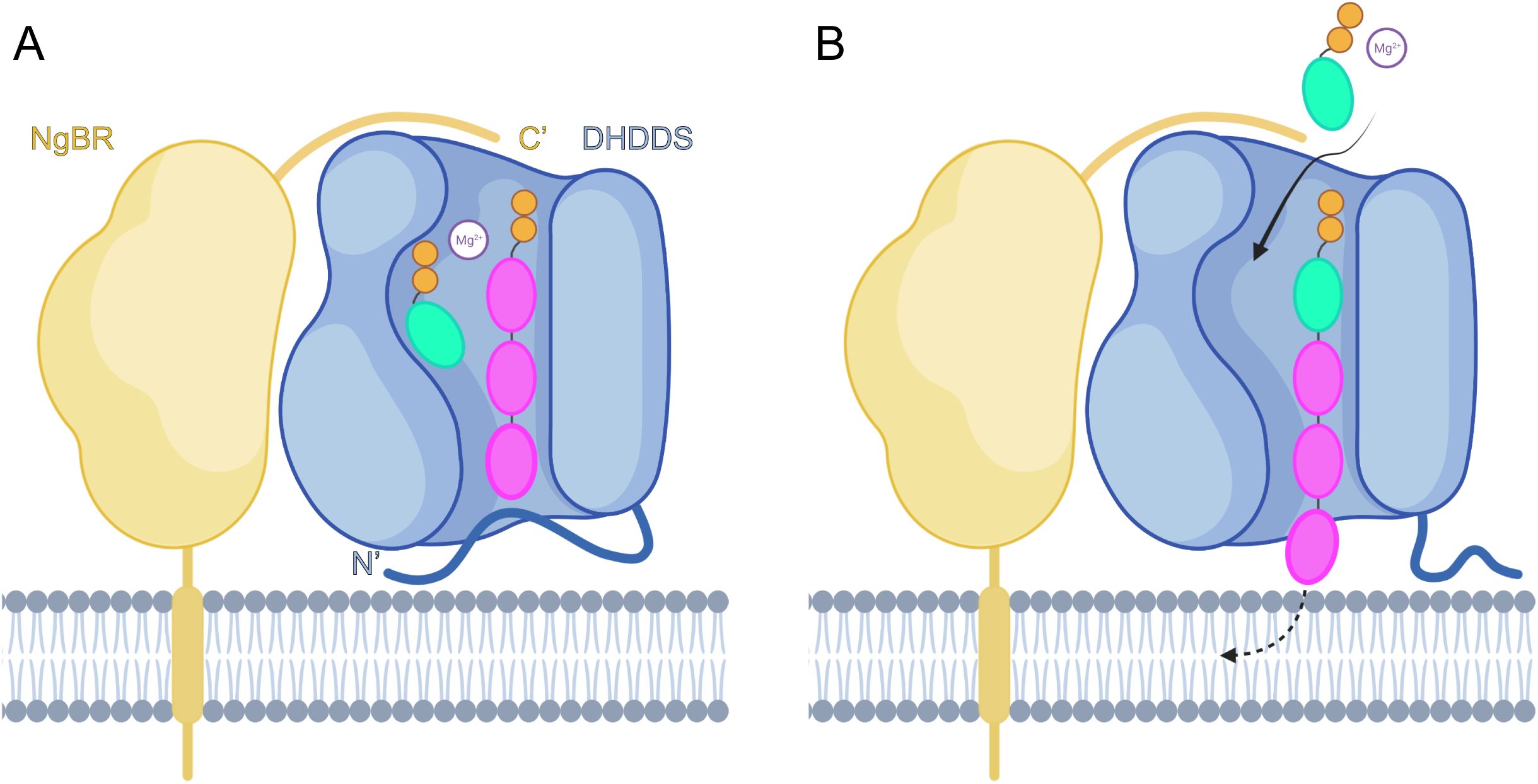
Proposed model for long-chain isoprenoids synthesis. (A) Prior to the initial condensation reaction, FPP (pink) and IPP (green) occupy the DHDDS active site. The active site outlet is occluded by the DHDDS N-terminus. (B) Following the condensation reaction, the N-terminus of DHDDS dislodges from the outlet, allowing extension of the elongating product to the vicinity of the ER membrane. Further product lengthening is enabled by its insertion into the bilayer. Created with BioRender.com.

A lingering question remains – what is the mechanism determining the final product length, preventing further condensations and leading to product release? Interestingly, in contrast with short- and medium-chain enzymes, in which product length is intrinsically determined by active site dimensions, long-chain *cis*-prenyltransferases and rubber synthases exhibit a population of products of varying lengths. This can reflect a tug-of-war mechanism, where the probability of product release is positively correlated with chain lengthening due to cumulative product-membrane interactions and its conformation within the lipid bilayer. Importantly, such a mechanism can account for the reduction in chain length resulting from disease-causing mutations at the pyrophosphate binding region,^22,37^ skewing the balance of forces in favor of the membrane, thus resulting in premature product release. Additional membrane properties, such as curvature, lipid composition, and fluidity may also play a role in product length determination.

In conclusion, our hybrid structural studies exposed a novel product outlet allowing direct and directional elongation of long-chain isoprenoids into adjacent membranes, providing a holistic view of h*cis*-PT catalysis and a structural basis for long-chain isoprenoids and natural rubber synthesis.

## Methods

### Protein expression and purification

*E. coli* T7 express competent cells were co-transformed with DHDDS (residues 1-333) and NgBR (residues 73-293) (sh*cis*-PT) or NgBRΔ167-175 (sh*cis*-PT_cryst_), grown in Terrific Broth medium at 37 °C until reaching OD_600nm_ = 0.6 and induced at 16 °C by adding 0.5 mM isopropyl β-D-1-thiogalactopyranoside (IPTG). Proteins were expressed at 16 °C for 16-20 hr, harvested by centrifugation (~5,700xg for 15 min), and then resuspended in a buffer containing 20 mM 4-(2-hydroxyethyl)-1-piperazineethanesulfonic acid (HEPES), pH 7.5, 150 mM NaCl, and 1 mM tris(2-carboxyethyl)phosphine (TCEP) and 0.02% (w/v) triton X-100, supplemented with 1 μg/ml DNase I and a protease inhibitor mixture. Resuspended cells were homogenized and disrupted in a microfluidizer. Soluble proteins were recovered by centrifugation at ~ 40,000xg for 45 min at 4 °C. Overexpressed proteins were purified on a HisTrap HP column, followed by purification on a Strep-Tactin column and TEV protease cleavage of the purification tags and TRX fusions. The reaction mixture was concentrated and loaded onto a Superdex-200 preparative size-exclusion column pre-equilibrated with 20 mM 4-(2-hydroxyethyl)-1-piperazineethanesulfonic acid (HEPES), pH 7.5, 150 mM NaCl, 1 mM tris(2-carboxyethyl)phosphine (TCEP). Purified proteins were flash-frozen in liquid nitrogen and stored at -80 ° C until use. Protein purity was >95%, as judged by SDS-PAGE.

### Crystallization and data collection

All crystallization trials were performed at 19 °C using the sitting drop vapor diffusion method. The sh*cis*-PT_cryst_-FsPP-IPP complex was crystallized by mixing a protein solution at ~16 mg/mL in the presence of 0.5 mM MgCl_2_, 0.75 mM FsPP and 1.5 mM IPP with 0.1 M NaCl, 0.1 M 2-(N-morpholino)ethanesulfonic acid (MES) pH 6.5, 33% w/v PEG 400. Data were collected at 100 K using wavelength of 0.976Å at the Diamond Light Source (DLS; Oxfordshire, United Kingdom) beamline I03 (https://doi.org/10.2210/pdb7PAX/pdb). The sh*cis*-PT_cryst_-GGsPP complex was crystallized by mixing a protein solution at ~14 mg/mL in the presence of 0.5 mM MgCl_2_, 1 mM FPP and 2 mM IsPP with 1 M LiSO_4_, 0.02 M Tris-HCl pH 8.5, 1.8% w/v PEG 8000. Crystals were cryoprotected with 20% glycerol and data were collected at 100 K using wavelength of 1.000Å at the European Synchrotron Radiation Facility (ESRF; Grenoble, France) beamline ID-30B (https://doi.org/10.2210/pdb7PAY/pdb). The sh*cis*-PT_cryst_-GGsPP-IsPP complex was crystallized by mixing a protein solution at ~12 mg/mL in the presence of 0.5 mM MgCl_2_, 1 mM FPP and 2 mM IsPP with 0.1 M KCl, 0.1 M MES pH 6.5, 36% w/v PEG 400. Data were collected at 100 K using wavelength of 1.000Å at the ESRF beamline ID-30B (https://doi.org/10.2210/pdb7PB0/pdb). The sh*cis*-PT_cryst_-GGPP-IsPP complex was crystallized by mixing a protein solution at ~4 mg/mL in the presence of 0.5 mM MgCl_2_, 1 mM GGPP and 2 mM IsPP with 0.07 M MES pH 6.6, 25% w/v PEG 400, 1.5 mM MgCl_2_, 4.5% v/v ethanol. Data were collected at 100 K using wavelength of 0.976Å at the ESRF beamline ID-23-1 (https://doi.org/10.2210/pdb7PB1/pdb). Integration, scaling and merging of the diffraction data were done with the XDS program.^38^ In the case of sh*cis*-PT_cryst_-GGPP-IsPP, a dataset with low completeness (~75%) at a resolution of 2.6Å was merged with a complete dataset (99.1%) at a resolution of 3.1Å using XDSCONV and XSCALE.^38^

### Structure determination and refinement

All the structures were solved by automated molecular replacement using Phaser^39^ with the structure of sh*cis*-PT_cryst_ -FPP (PDB 6Z1N) as a search model. Iterative model building and refinement were carried out in PHENIX^40^ with manual adjustments using COOT.^41^ Ramachandran analysis was performed using MolProbity.^42^ Data collection and refinement statistics are presented in Table 1. Structural illustrations were prepared with UCSF Chimera (https://www.cgl.ucsf.edu/chimera). Atomic coordinates and structure factors for the structures of sh*cis*-PT_cryst_ have been deposited in the Protein Data Bank under accession codes 7PAX (in complex with Mg^2+^, FsPP and IPP), 7PAY (in complex with Mg^2+^ and GGsPP), 7PB0 (in complex with Mg^2+^, GGsPP and IsPP), and 7PB1 (in complex with Mg^2+^, GGPP and IsPP).

### HDX-MS

For peptide identification, sh*cis*-PT was diluted to 2uM concentration in 20 mM HEPES, pH 7.5, 150 mM NaCl, 1 mM EDTA, 1 mM TCEP and subsequently mixed with equal volume (50 μL + 50 μL) of 1 M glycine-HCl, pH 2.3. Mixture was frozen in liquid nitrogen. Prior to the analysis, the sample was quickly thawed and injected onto an LC system coupled to an ESI source of 15T FT-ICR mass spectrometer (solariX XR, Bruker Daltonics, Bremen, Germany) operating in data dependent mode, where each full scan was followed by six MSMS scans (CID) of the top most abundant ions. Repeat count was set to 2, exclusion was set to 10 ppm and 30 sec duration. Instrument was calibrated externally using sodium TFA clusters resulting in sub-ppm mass error. The LC method started by an online digestion on the custom-made pepsin column (bed volume 66 µL).^43^ Generated peptides were trapped and desalted on a VanGuard Pre-column (ACQUITY UPLC BEH C18, 130Å, 1.7 μm, 2.1 mm × 5 mm, Waters, Milford, MA, USA). The digestion and desalting (total time 3 min) were driven by 0.4% formic acid (FA) in water pumped by 1260 Infinity II Quaternary pump (Agilent Technologies, Waldbronn, Germany) at a flow rate of 100 μL·min^−1^. Desalted peptides were then separated on an analytical column (ACQUITY UPLC BEH C18, 130 Å, 1.7 μm, 1 mm × 100 mm, Waters, Milford, MA, USA) using linear gradient (5%–45% B in 7 min) followed by a quick step to 99% B lasting 5 min, where the solvent A was 0.1% FA /2% acetonitrile (ACN) in water, B was 0.1% FA /98% ACN in water. The gradient was delivered by the 1290 Infinity II LC System (Agilent Technologies, Waldbronn, Germany) at a flow rate 40 μL·min^−1^. LC-MS/MS analysis for peptide identification was followed by MASCOT database searching (version 2.7, Matrix Science, London, United Kingdom) against a custom-built database combining crap database, sequence of used acidic protease and the sequences of sh*cis*-PT subunits. MSTools was used to plot the identified peptides against the protein sequence.^44^

Backbone amide HDX was initiated by a 10-fold dilution of the protein solution into D_2_O-based solution containing 20 mM HEPES, pD 7.4, 150 mM NaCl, 1 mM EDTA, 1 mM TCEP. The exchange was followed for 20 sec, 1 min, 3 min, 10 min, 30min, 1 hr and 3 hr at 20 °C and quenched by the addition of 0.5 M glycine-HCl, pH 2.3 in a 1:1 ratio and flash freezing in liquid nitrogen. The labelling experiment was done in duplicate. LC-MS analysis of deuterated samples was started by rapid thawing of the sample and injection onto the custom-made pepsin column (bed volume 66 µL). Following LC-method was identical to that used for peptide identification and mentioned above. ^43^ 15T FT-ICR mass spectrometer was operating in broad-band MS mode. Acquired data were exported using DataAnalysis v. 5.0 (Bruker Daltonics, Bremen, Germany), processed by the in-house designed Deutex software and handled as described previously.^45,46^ Fully deuterated control was prepared as described previously^46^ and correction for back-exchange was applied.^32^

### Enzyme kinetics

The activity of purified sh*cis*-PT was measured using a radioligand-based assay. ^12,26,47,48^ 0.1 μM of purified sh*cis*-PT were mixed with MANT-O-GPP and [^14^C]-IPP to initiate the reaction in buffer composed of 25 mM Tris-HCl, pH 7.5, 150 mM NaCl, 10 mM β-mercaptoethanol, 0.02% Triton-X100, 0.5 mM MgCl_2_ at 30 °C. 15 mM EDTA (final concentration) were added to quench the reaction and 500 µL of water-saturated 1-butanol was added to extract the reaction products by thorough vortexing. Initial rates were measured by quenching the reaction at 10% or lower substrate consumption. The products, encompassing ^14^C, were quantitated using a scintillation counter. The *K*_M_ value of MANT-O-GPP was determined by varying [MANT-O-GPP] while holding [IPP] constant at 100 μM. Kinetic constants were obtained by fitting the data to the Michaelis-Menten equation using Origin 7.0 (OriginLab, USA).

### Fluorescence spectroscopy

MANT-O-GPP was synthesized as previously described. All fluorescence experiments were performed in triplicates using a Jasco RF-8500 spectrofluorometer in fluorescence buffer (20 mM Tris-HCl, pH 7.5, 150 mM NaCl, 10 mM β-mercaptoethanol, 0.5 mM MgCl_2_). FRET experiments between sh*cis*-PT and MANT-O-GPP were performed using 1 μM of purified sh*cis*-PT (WT or W3L) preincubated with 0-16 μM MANT-O-GPP. The competition with FPP was performed by preincubation with varying [FPP] (0-100 μM) while holding [MANT-O-GPP] constant at 10 μM. To obtain the dose-response curves, the *F*_420_/*F*_340_ ratio was calculated for each fluorescence spectrum and the *F*_420_/*F*_340_ ratio of the apo protein was subtracted. The activity of sh*cis*-PT in the presence of azolectin nanodiscs was measured by using 0.1 μM of purified sh*cis*-PT preincubated with the indicated ratio of nanodiscs in the presence of 2 μM MANT-O-GPP and 40 mM IPP. Experiments with fluorescent nanodiscs were carried out under the same conditions using a constant concentration of 0.2 μM nanodiscs. Data were fit using Prism GraphPad 9.0.1 (GraphPad Software, USA).

### Nanodiscs preparation

Membrane scaffold protein 1E3D1 (MSP1E3D1) was overexpressed and purified as previously described.^49^ Briefly, hexahistidine-tagged MSP1E3D1 was overexpressed in *E. coli* and purified using immobilized metal affinity chromatography followed by tag cleavage with TEV protease and size-exclusion chromatography in nanodiscs buffer (20 mM Hepes, pH 7.5, 150 mM NaCl). Nanodiscs were prepared by mixing 0.2 mM MSP1E3D1 in nanodiscs buffer supplemented with 10 mM azolectin in the presence or absence of 1% mol/mol PE-NBD in 50 mM cholate. Nanodiscs buffer was added to achieve a final cholate concentration of 20 mM and the nanodiscs assembly mixture was incubated for 1 hr at room temperature. Next, 1 mg/μl bio-beads were added in two steps, where half of the bio-beads were incubated for 1 hr and the other incubated for 3 hr at room temperature. Finally, the reaction volume was centrifuged for 5 minutes at 21,000xg and loaded onto a Superdex-200 preparative size-exclusion column pre-washed with nanodiscs buffer.

## Supporting information

Figures depicting the crystal packing and heterotetrameric assembly (Figure S1), superposition between the complex bound to FsPP-IPP, FPP or IPP (Figure S2) and sequence coverage of the HDX-MS experiments (Figure S3).

## Acknowledgments

We acknowledge the European Synchrotron Radiation Facility for provision of synchrotron radiation facilities and we would like to thank Dr. Montserrat Soler-López and Dr. Christoph Mueller-Dieckmann for assistance in using beamlines ID-23-1 and ID-30B, respectively. We also acknowledge the Diamond Light Source for provision of synchrotron radiation facilities and we would like to thank Ms. Felicity Bertram for assistance in using beamline I03. This work was supported by the Israel Science Foundation grants 1721/16 and 1653/21 (Y.H.), the Israel Cancer Research Fund grants 01214 (Y.H.) and 19202 (M.G.), the Israel Cancer Association grant 20200037 (M.G. and Y.H.), and from the German□Israeli Foundation for Scientific Research and Development grant I□2425□418.13/2016 (Y.H.). Support also came from the I□CORE Program of the Planning and Budgeting Committee and The Israel Science Foundation grant 1775/12 (Y.H.) and the Claire and Amedee Maratier Institute for the Study of Blindness and Visual Disorders, Sackler Faculty of Medicine, Tel-Aviv University (Y.H. and M.G.). Access to MS installation was funded by the EU Horizon 2020 grant EU_FT–ICR_MS project number 731077 and by CIISB LM2018127. P.M. and P.V. support from MEYS CZ funds CZ.1.05/1.1.00/02.0109 is gratefully acknowledged.

## Notes

### Competing Interest Statement

The authors have declared no competing interest.

