## Supplemental Figures 1-3 for "Structural basis for long-chain isoprenoids synthesis by *cis*-prenyltransferases"

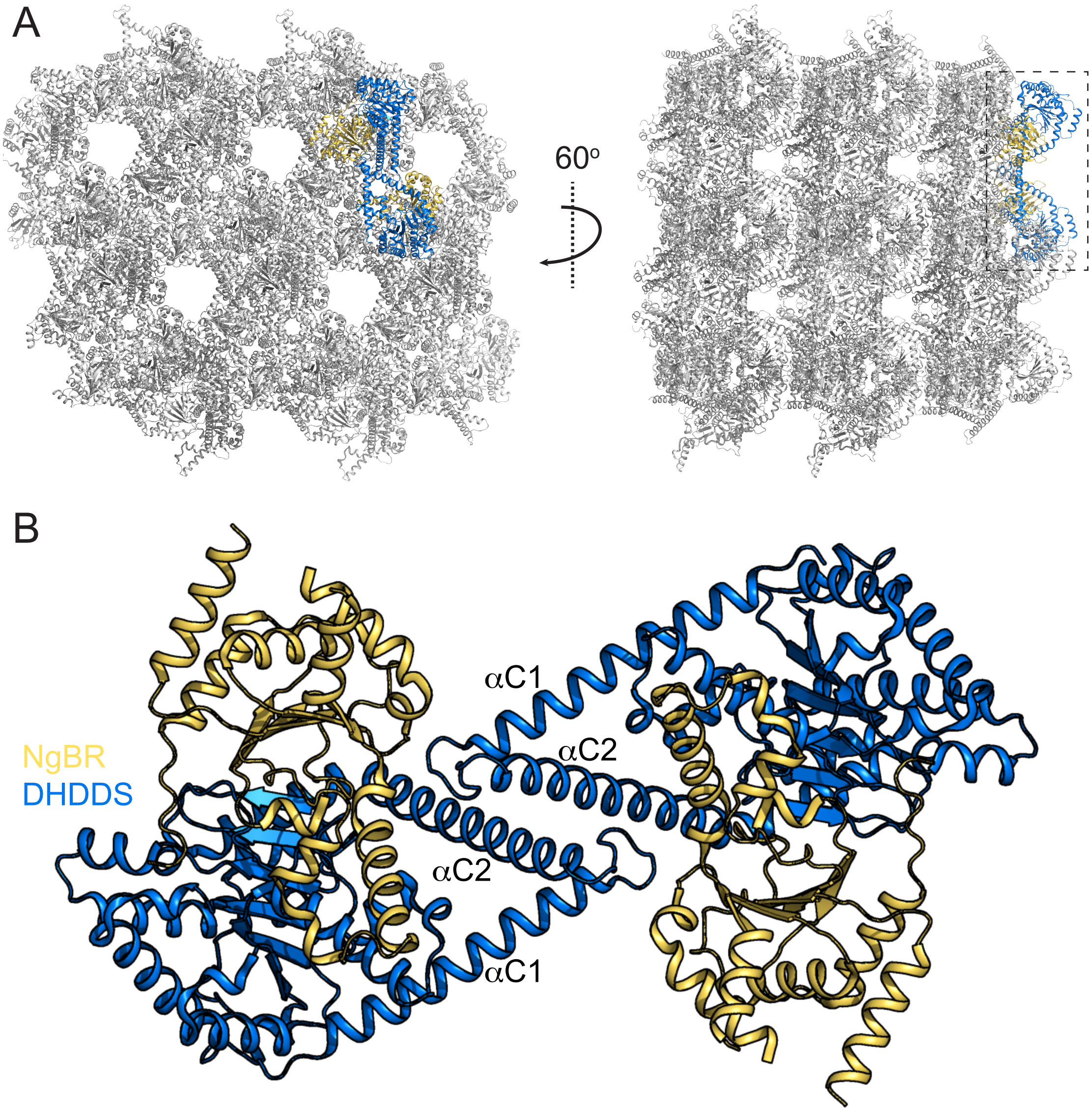


**Figure S1.** Crystal packing and heterotetrameric assembly of sh*cis*-PT$_{cryst}$. (A) Crystal packing of sh*cis*-PT$_{cryst}$ (PDB 6Z1N) in the R32:H space group is shown in two projections in cartoon representation. A single heterotetramer, formed by adjacent asymmetric units, is framed by a dashed rectangle (right) and highlighted in color as in B. (B) Cartoon representation of the sh*cis*-PT$_{cryst}$ heterotetramer. DHDDS and NgBR are colored blue and yellow, respectively. αC1 and αC2, forming the dimerization surface between two DHDDS-NgBR heterodimers, are indicated.

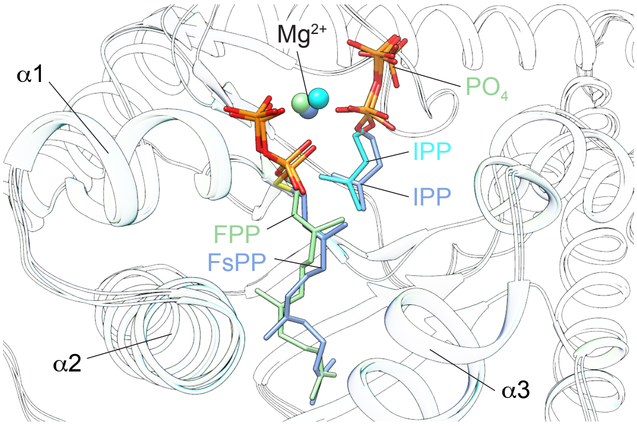


**Figure S2.** Superposition of the structures of sh*cis*-PT in complex with FsPP-IPP (light blue), IPP (cyan) or FPP (green) in cartoon representation. The substrates are shown as sticks and the $Mg^{2+}$ ions are shown as spheres.

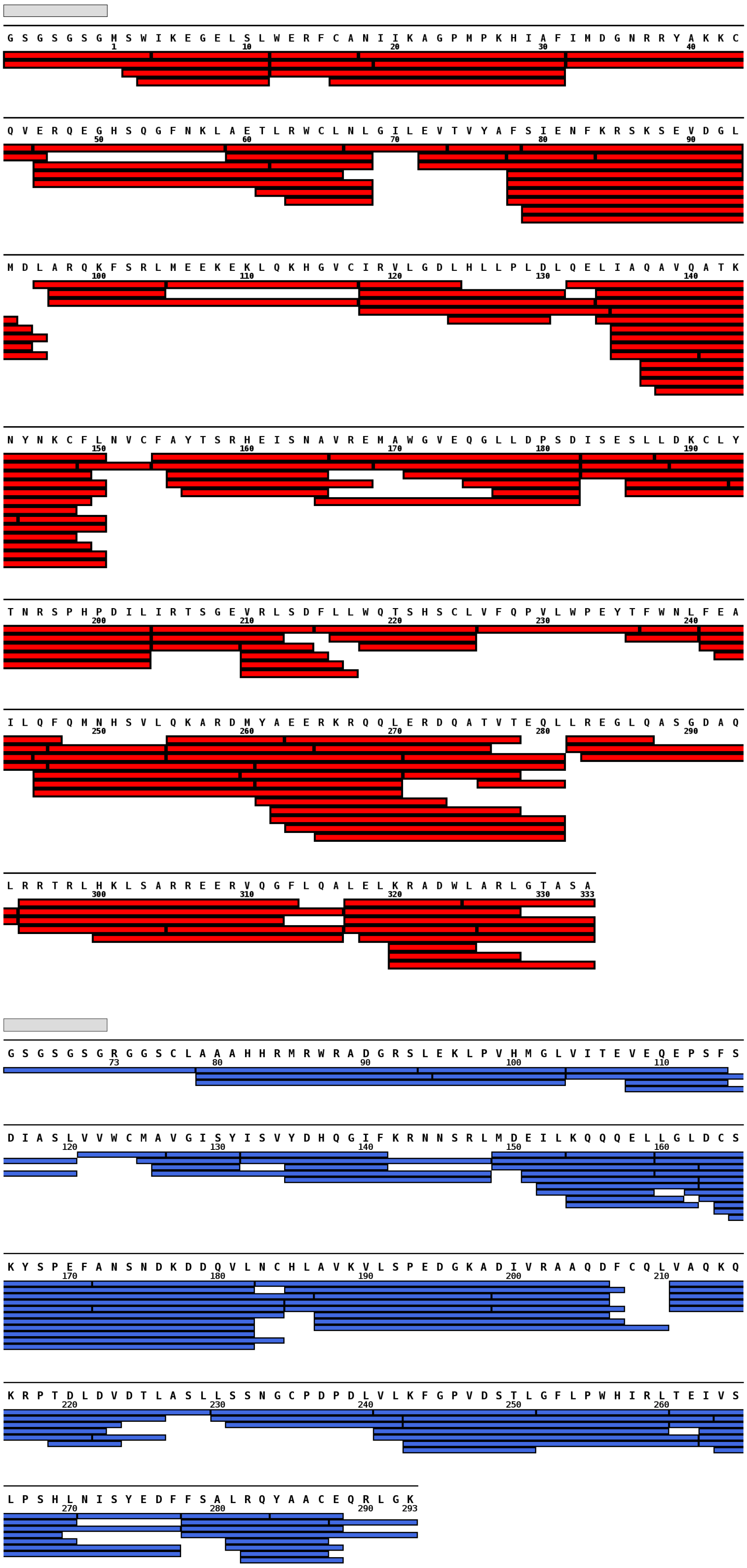


**Figure S3. Hydrogen-deuterium exchange mass-spectrometry (HDX-MS) sequence coverage.** Peptic peptides are represented as red or blue bars under the sequences of DHDDS or NgBR, respectively. A remnant of the expression tag following TEV digestion is indicated with a gray bar above each sequence.
